# Single dose alum adjuvanted RBD protein vaccine provides protection against homologous challenge with SARS-CoV-2 Washington strain and heterologous rechallenge with Delta and Omicron BA.5 variants in K18 hACE2 mouse model

**DOI:** 10.1101/2025.09.18.677166

**Authors:** Rajesh Thippeshappa, Viraj Kulkarni, Varun Dwivedi, Jennifer McMillian, Marco Argonza, Joshua Castro, Oscar Rodriguez, Billie Maingot, Michal Gazi, Reagan Meredith, Deepak Kaushal, Ricardo Carrion

## Abstract

Most COVID-19 vaccination strategies require at least two doses-a primary vaccine followed by a booster dose with the updated variant-specific vaccine. However, vaccine and booster dose hesitancy make it challenging to administer multiple doses of vaccines in the unvaccinated and vaccinated populations, respectively. Thus, it is important to determine if vaccinated individuals exposed to SARS-CoV-2 infection develop immune responses that may protect them against emerging variants of concern (VoCs). We have developed mouse models to understand the protective efficacy of an RBD-based vaccine against challenge and rechallenge with SARS-CoV-2 VoCs. Mice were vaccinated with RBD protein vaccine formulated in 2% Alhydrogel (alum) adjuvant by subcutaneous route. To determine the efficacy of RBD vaccine, mice were challenged approximately 4 weeks post-vaccination with SARS-CoV-2 variants by intranasal route. To determine the efficacy of RBD vaccine against SARS-CoV-2 rechallenge, mice were rechallenged with SARS-CoV-2 Delta and Omicron BA.5 variants at day 14 post SARS-CoV-2 Washington (WA) strain challenge. Our data suggest that single dose alum adjuvanted RBD protein from Wuhan strain provides protection against homologous challenge with SARS-CoV-2 WA strain but failed to provide protection against heterologous challenge with Delta and Omicron BA.5 variants. Interestingly, vaccinated mice that survived homologous challenge with the WA strain showed protection against heterologous rechallenge with Delta and omicron BA.5 variants. Furthermore, infectious viral loads of Delta and Omicron BA.5 were not detected in the lung tissues collected from the rechallenged mice at 3 days post-rechallenge. The data suggest that a single dose RBD vaccine from the ancestral Wuhan strain together with survival from WA strain challenge induces protective immune responses against Delta and Omicron BA.5 variants rechallenge. These mouse models will be useful to determine the immune responses that correlate with protection against challenge and rechallenge with SARS-CoV-2 VoCs.

## INTRODUCTION

COVID-19, characterized by severe pneumonia and acute respiratory distress syndrome, is caused by a recently identified coronavirus named SARS-CoV-2. Although there are several approved vaccines, global vaccination rates are low due to inaccessibility (1) (WHO COVID19 dashboard). The two widely used mRNA-based vaccines are expensive and require ultra-low temperatures for storage and transportation (2). This presents a challenge in administering these vaccines in resource-limited low and middle-income countries (LMICs). Therefore, there is a great need for developing vaccines with storage properties that will make them implementable in low-resource settings. In this regard, protein subunit vaccines that are easy to manufacture and stable at ambient temperature may facilitate improved vaccination rates in LMICs. Subunit vaccines have been developed against infectious diseases such as hepatitis B, diphtheria, tetanus, and shingles (3, 4). However, there is only one FDA-approved subunit-based vaccine against SARS-CoV-2 (Novavax). This vaccine is based on recombinant full-length spike protein formulated in a saponin-derived Matrix-M adjuvant. Although full-length spike protein is immunogenic and an attractive target for vaccine development, there are also efforts to develop subunit-based vaccines based on smaller fragments of the spike protein containing the receptor binding domain (RBD) (5, 6). This is because the attachment of SARS-CoV-2 to its receptor angiotensin-converting enzyme 2 (ACE2) is mediated through RBD of spike protein (7, 8), and antibodies against RBD can neutralize SARS-CoV-2 by blocking interaction with ACE2. Additionally, due to its thermal stability, low-cost manufacturing, and scalability, RBD presents an attractive target for developing subunit vaccine candidates (9–13). Adjuvanted RBD-based vaccines have been reported to induce robust neutralizing antibody responses and protection in SARS-CoV-2 challenge studies (13–24). Furthermore, RBD-based vaccines containing both alum and CpG adjuvants has received emergency use authorization in India and Indonesia (25).

Our initial efforts focused on developing “Prime-Boost” vaccination approach: priming with recombinant vectors expressing receptor binding domain protein of SARS-CoV-2 spike protein and boosting with recombinant RBD protein-based subunit vaccine. Prime-Boost vaccination approach has been used to induce both humoral and cell mediated immune responses to many pathogens (26–29). For priming, we focused our efforts to develop recombinant BCG expressing SARS-CoV-2 immunological targets of interest. This is because, epidemiological studies as well as clinical trials suggest non-specific beneficial effects of BCG vaccination on respiratory tract infections caused by unrelated pathogens (30, 31). The molecular mechanisms contributing to nonspecific beneficial effects of BCG vaccination suggest long-term activation and reprograming of innate immune cells, a phenomenon termed trained immunity (32). Studies suggest that epigenetic and metabolic programing of monocytes by BCG vaccination results in enhanced expression of pro-inflammatory cytokines during subsequent viral infections (33, 34). Therefore, it is thought that induction of trained immunity by BCG vaccination could provide significant protection against viral infections (30, 35). Thus, we hypothesized that priming with rBCG-RBD and boosting with adjuvanted RBD protein subunit vaccine will induce both specific and non-specific beneficial immune responses, thereby conferring better protection against SARS-CoV-2 infection. We determined immunogenicity and protective efficacy of rBCG-RBD prime and alum adjuvanted recombinant RBD protein boost approach in K18 hACE2 transgenic mouse model, which have been shown to be highly susceptible to SARS-CoV-2 Isolate 2019-nCoV**/**USA-WA1/2020 (referred to as WA strain) infection (36). Our data suggest that there was no significant difference in the protection offered by RBD protein vaccine alone compared to rBCG-RBD prime and RBD protein boost approach. Consequently, we focused our efforts to determine the protective efficacy of alum adjuvanted RBD protein vaccine against heterologous challenge with SARS-CoV-2 variants of concern (VoCs). Interestingly, single dose RBD vaccine did not provide protection against heterologous challenge with SARS-CoV-2 Delta and Omicron BA.5 variants.

Many individuals vaccinated with either ancestral or recent COVID-19 vaccines may not take updated boosters. Because they have probably been exposed to one or more of SARS-CoV-2 variants, it is important to determine if ancestral or recent COVID-19 vaccines combined with virus exposure induces protective immune responses against future variants of concern (VoCs). We have developed a mouse model to determine if a single dose RBD vaccine combined with virus exposure would induce protective immune responses against SARS-CoV-2 VoCs. Interestingly, single dose RBD vaccinated mice that survived homologous challenge with WA strain showed protection against heterologous challenge with Delta and Omicron BA.5 variants. Data suggest that single-dose vaccine combined with virus infection may induce multiple protective immune responses, that may provide protection against future SARS-CoV-2 VoCs. Understanding these immune responses and determining if they correlate with protection against SARS-CoV-2 challenge and rechallenge will be useful in the development of novel and updated vaccines against recent SARS-CoV-2 VoCs.

## MATERIALS AND METHODS

### Cell lines

Vero E6 and Vero E6 expressing human angiotensin and TMPRSS2 (Vero-AT) cells were cultured in Dulbecco’s modified Eagle’s medium (DMEM) supplemented with 10% heat-inactivated fetal bovine serum (HI-FBS), 2 mM glutamine, 100 U of penicillin per ml and 100 μg of streptomycin per ml (p/s) (DMEM complete).

### Virus stocks

SARS-CoV-2, USA-WA1/2020 strain (BEI Resources NR-52281), SARS-CoV-2 delta variant (B.1.617.2) (NR-55611), and SARS-CoV-2 omicron BA.5 variant (B.1.617.2) (NR-52281) were obtained from The Biological and Emerging Infections Resources Program. A 6th passage (P6) of SARS-CoV-2 WA strain and a 5th passage (P5) of SARS-CoV-2 delta variant were generated by infecting Vero E6 cells for 72 hours (h). A 3rd passage (P3) of SARS-CoV-2 omicron BA.5 was generated by infecting Vero E6 expressing hACE2 and TMPRSS2 cells (Vero-AT) for 72 hours (h). At 72 h post infection, culture supernatants were collected, clarified, and aliquots were stored at −80°C. Viral titers were determined by plaque assay.

### Recombinant BCG (rBCG) constructs

steps described below were performed to generate rBCG constructs

### Preparation of cDNA

SARS-CoV-2 RNA was reverse transcribed using random hexamer primer or oligo(dT) primers using RevertAid RT kit (Thermo Fisher Scientific) according to manufacture instructions. cDNA was then amplified with below primer sets for amplification of N and RBD gene fragments using superfi II green PCR mix.

### Amplification of N gene

SD-CoV-N-F: ACG TAG AAG GAG AAG TAC CGA TGT CTG ATA ATG GAC CCC AAA ATC AGC and PMV-CoV-N-R: GTA CGC TAG TTA ACT TTA GGC CTG AGT TGA GTC AGC ACT GCT CA. PCR conditions: were 98°C for 30 sec, then 98°C for 10 s, 61°C for 10 s, and 72°C for 10 min for 35 cycles; and then 72°C for 5 min.

### Amplification of RBD-1

SD-RBD-319F: ACG TAG AAG GAG AAG TAC CGA TGA GAG TCC AAC CAA CAG AAT CTA TTG and RBD-541R: GAT CGT ACG CTA GTT AAC TCT AGA AAT TGA CAC ATT TGT TTT TAA CCA

### Amplification of RBD-2

SD-RBD-319F: ACG TAG AAG GAG AAG TAC CGA TGA GAG TCC AAC CAA CAG AAT CTA TTG and RBD-591R: GAT CGT ACG CTA GTT AAC TCT AAG AAC ATG GTG TAA TGT CAA GAA TCT C

### Vector Backbone PCR for insertion of N gene

PMV-306hsp vector was amplified with PMV-SD-R: CGG TAC TTC TCC TTC TAC GTC GAC ATC GAT AAG CTT G and CoV-N-PMV-F: CoV-N-PMV-F. PCR conditions: were 98°C for 30 sec, then 98°C for 10 s, 60°C for 10 s, and 72°C for 10 min for 35 cycles; and then 72°C for 5 min.

### Vector Backbone PCR for insertion of RBD genes

PMV-306hsp vector was amplified with PMV-SD-R: CGG TAC TTC TCC TTC TAC GTC GAC ATC GAT AAG CTT G and PMV-FWD AGT TAA CTA GCG TAC GAT CGA C primers. PCR conditions: were 98°C for 30 sec, then 98°C for 10 s, 60°C for 10 s, and 72°C for 10 min for 35 cycles; and then 72°C for 5 min.

#### NEBuilder HiFi DNA assembly

Vector PCR product and insert PCR product were mixed with a minimum of 1:3 ratio in a 20 μl reaction volume containing 10 μl of HiFi assembly mix. After 1 hour of incubation at 50 °C, 2 to 5 μl of reaction mix was used for transformation of competent cells (New England BioLabs). Miniprep plasmids were isolated using QIAprep spin miniprep kit (Qiagen).

### Electroporation of competent BCG

Competent BCG Danish 1331 strain cell aliquots (100 µl) were thawed on ice and 1-5 μg was added and mixed by gentle pipetting. The mix was transferred to a pre-chilled GenePulser electroporation cuvette (0.2-cm electrode gap, Bio-Rad). A single pulse was applied with a Gene Pulser Xcell PC apparatus (Bio-Rad, USA) which was set at 2500 V, resistance 200 Ω, capacitance 25 µF. Immediately after the pulse the cells were mixed with 5 ml of 7H9 MB medium supplemented with 0.05% Tween-80 and 10% ADC in 50ml sterile culture tubes and recovered at 37°C for 24 hours. Finally, the cells were plated out onto 7H11 Middlebrook (10% ADC) agar plates with 20 μg/ml kanamycin and incubated for 15-30 days at 37°C until the colonies appeared

### Colony PCR

Colonies were picked from the 7H11 agar plates and inoculated into a PCR tube containing 10 µl sterile water. Superfi II green PCR mix containing gene specific primers was then added to the PCR tube. Same PCR conditions described for amplification of N and RBD region of S gene were used to identify positive colonies.

### Growth of rBCG

rBCG stocks were generated by growing individual colonies in Middlebrook 7H9 media (10% ADC) containing 20 µg/ml kanamycin at 37°C in a shaker incubator. Cultures in exponential phase (OD600=0.8) were washed twice in ice cold PBS and cell pellet was resuspended in PBS supplemented with 50% glycerol. rBCG stocks were frozen at −80°C.

### Western blot

Colonies that were PCR positive for gene of interest were grown in 7H9 media (10% ADC) containing 20 µg/ml kanamycin to confirm expression of protein by western blot. 10ml of culture was pelleted at 3000 rpm for 7 minutes. Pellet was resuspended in 500 µl cold DPBS with 1X protease inhibitor and transferred to Zirconian tubes. Bacterial lysate was prepared using mini BeadBeater set at 4.5 m/s, 30seconds, for a total of 5 times. Tubes were centrifuged at 14000 rpm for 20 minutes. Supernatant was collected at stored at −20°C.

Mini-PROTEAN TGX precast proteins gels (4-20%) were used in western blot. Sample buffer (4X) and bacterial lysate were mixed 1:3 and boiled for 10 min. 50 µl sample was loaded into each well and run at 100 V for 90 minutes. Gel was then transferred to PVDF membrane using semidry transfer method. Membrane was washed once in 1x TBS with 0.05% tween 20 (TBST) and blocked for 1 hour at room temperate on shaker with 5% non-fat dry milk in TBST. Membrane was washed 3 times with TBST before incubating with primary antibody in 1 % blocking solution. Membrane was washed again 3 times in TBST and incubated with secondary antibody (1:5000) in 1 % blocking solution. After 3 washes with TBST, membrane was added with supersignal west femto maximum sensitivity substrate. Antibody to N protein was a generous gift from Dr. Luis Martinez, Texas Biomedical Research Institute. Antibody to RBD protein was purchased from R&D systems (Cat # MAB10540-100).

### Mice immunization and infection

#### Ethics statement

All mouse experiments were performed according to ethical guidelines by the Institutional Animal Care and Use Committee (IACUC) of Texas Biomedical Research Institute.

#### Mice

Transgenic K18hACE2 [B6.Cg-Tg(K18-ACE2)2Prlmn/J] mice, female, 4- to 6-week-old, were obtained from Jackson laboratories. Mice were quarantined for a week before the start of the experiment. Mice were maintained 5/cage within individual micro-isolator cages within a HEPA filtered rack, and cages were changed weekly. Mice were housed in the ABSL2 vivarium during the immunogenicity phase of the study and transferred to ABSL3 lab for SARS-CoV-2 infection.

### Vaccine preparation and administration

#### rBCG

Recombinant BCG was thawed and washed 2 times in ice cold PBS and resuspended in PBS for desired concentration. Mice were injected subcutaneously with either 1×10^8^ cfu of wildtype BCG or recombinant BCG. Unvaccinated mice received PBS alone.

#### Protein Immunogens

Spike Glycoprotein Receptor Binding Domain (RBD) from SARS-Related Coronavirus 2, Wuhan-Hu-1 with C-Terminal Histidine Tag, recombinant from HEK293T Cells was obtained from BEI Resources (NR-52946). SARS-CoV-2 recombinant nucleocapsid (N) protein was commercially purchased from Thermofisher Scientific (Cat # RP-87665). For protein vaccination, 5 μg of recombinant purified N or RBD protein formulated in Alhydrogel adjuvant 2% (alum) (InvivoGen) was given subcutaneously to each mouse. Unvaccinated mice received adjuvant alone. Mice were bled every 14 days to collect serum to determine antibody titer.

### Mice Infection and tissue processing

Mice were challenged intranasally with 1×10^3^ pfu of SARS-CoV-2 WA strain and Delta variants. For challenge with Omicron BA.5 variant, 1×10^5^ pfu was used. More than 20% loss of initial body weight was considered an experimental endpoint and mice were humanely euthanized to collect blood and tissues. For viral load determination, lung lobes were collected, harvested, and homogenized in 1 ml of sterile PBS using a Precelly’s tissue homogenizer (Bertin Instruments, Rockville, MD). Tissue homogenates were centrifuged at 8,000 x *g* for 15 min at 4°C and supernatants were collected and stored at −80°C.

#### Viral load quantitation by plaque assay

Virus stock titers and viral titers in tissue homogenates were determined by plaque assay described in detail (37, 38). Briefly, confluent monolayers of Vero E6 or Vero-AT cells (24-well plate format, 2X10^5^ cells/well) were infected with 100 μl of ten-fold serial dilutions of tissue homogenates diluted in DMEM medium for 1 h at 37°C, 5% CO_2_, with intermittent rocking every 15 min. After viral adsorption, cells were overlaid with 800 μl of post-infection media [DMEM + 2% FBS + PSG] containing 1% low melting agar and incubated for 48 h at 37°C, 5% CO_2_. Subsequently, cells were fixed overnight with 10% formaldehyde solution. For immunostaining, cells were washed three times with PBS, and permeabilized with 0.5% Triton X-100 for 10 min at room temperature. Cells were immunostained with a SARS-CoV/SARS-CoV-2 nucleocapsid (N) protein cross-reactive monoclonal antibody (mAb, 1C7C7, Cat# ZMS1075, Sigma-Aldrich, Saint Louis, MO) diluted (1 μg/ml) in 1% Bovine Serum Albumin (BSA) for 1 h at 37°C. After incubation with the primary 1C7C7 mAb, cells were washed three times with PBS and developed with the Vectastain ABC and DAB Peroxidase Substrate kits (Vector 580 Laboratory, Inc., CA, USA) following manufacturers’ instructions. Viral plaques were counted, and viral titers were calculated as PFU/ml.

#### Viral RNA loads by qRT-PCR assay

Viral load in tissue homogenates was determined by qRT-PCR. For virus inactivation, 100-250 µL of media was mixed with 750 µL of Trizol-LS reagent. Total RNA was isolated using NucleoMag Pathogen kit (Macherey-Nagel) in combination with KingFisher Flex instrument. Addition of MS2 phage to all samples was used as an internal control for the RNA extraction and qRT-PCR efficiency. (MS2-F: GGG TTT CCG TCT TGC TCG TA, MS2-R: TCG TGC TTT TCG CTG AAG AA, MS2-Probe: VIC-CGC TCG AGA ACG CAA). The genomic qRT-PCR for <u>SARS-CoV-2</u> was adapted from a CDC developed assay and is targeting a region of the N gene (N1-F: GAC CCC AAA ATC AGC GAA AT, N1-R: TCT GGT TAC TGC CAG TTG AAT CTG, N1-Probe: FAM-ACC CCG CAT TAC GTT TGG TGG ACC). Master Mix: TaqPath™ 1-Step RT-qPCR Master Mix, CG Thermo (Cat No. A15299). Cycling parameters: 25 °C 2 minutes, 50 °C 15 minutes, 95 °C 2 minutes, Amplification 45 x 95 °C 3 seconds, 60 °C 30 seconds. Analysis of tissue homogenates for RNA was performed on two replicates of each sample, and the mean is reported as GE/ μg of RNA.

#### ELISA

The antibody titer against SARS-CoV-2 RBD and N protein were measured by an enzyme-linked immunosorbent assay (ELISA) using recombinant RBD (Sino Biological) and N protein (Thermofisher Scientific) as the capture antigen. Each well of a 96-well microtiter plate (Corning, catalog # 2592) was coated with 100 ng of protein in 100 μl carbonate-bicarbonate buffer (pH 9.6) and incubated overnight at 4° C. Next day, plate was washed twice with 1X PBS containing 0.05% tween 20 (PBST) followed by blocking with 5% non-fat dry milk in PBST at room temperature for 2 hours. Two-fold serially diluted serum from vaccinated and control mice were added to duplicate wells and incubated at 37°C for 1 hour. After 5 washes with 1X PBST, 100 μl of Goat anti-mouse IgG peroxidase conjugate antibody (1:5000 dilution in blocking buffer) was added to each well and incubated at 37°Cfor 1 hour. Plate was washed 5 times with 1X PBST. SureBlue TMB peroxidase substrate (100 μl) was then added to each well and incubated at room temperature for 10 minutes. Reaction was stopped by adding 100 μl TMB stop solution, and the plate was immediately read at 450nm. The endpoint ELISA titer of binding antibodies was defined as the reciprocal of the serum dilution that resulted in positive optical density (OD) reading, which is at least two times the mean OD reading with no serum control wells. The detection limit of ELISA was considered to be the starting dilution (1:100) of the test serum.

#### Neutralization assay

A plaque reduction microneutralization (PRMNT) assay was used to evaluate the ability of the serum samples to neutralize SARS-CoV-2 (37, 38). Briefly, Vero E6 or Vero AT cells were seeded onto 96-well plates. The following day, serum samples (in triplicates) were serially diluted (2-fold) in DMEM supplemented with penicillin, streptomycin and L-glutamine. SARS-CoV-2 was added to the serially diluted serum samples at a multiplicity of infection (MOI) of 0.01 (100 to 200 PFU) per well. The antibody-virus mixture was incubated at 37°C for 1 h and then absorbed on cells for 1 h at 37°C. After viral absorption, the antibody-virus mixture was replaced with DMEM supplemented with penicillin, streptomycin and L-glutamine containing 1% Avicel. The plates were incubated at 37°C for 24 h. Following incubation, the plates were fixed in 10% formalin for 24 h. Cell monolayers were immunostained using the 1C7C7 mAb against the viral N protein and developed using an the Vectastain ABC and DAB Peroxidase Substrate kits. Viral plaques were scanned using the CTL ImmunoSpot plate reader and formed plaques counted by an automated counting software (Cellular Technology Limited, Cleveland, OH, USA).

#### Statistical Analysis Methods

Differences between survival curves for the unvaccinated and treatment groups were analyzed using the log-rank test. Proportions of surviving animals of the treatment groups compared to the unvaccinated groups was examined using Fisher’s Exact test. Differences in viral titers between vaccine and adjuvant groups were analyzed with Welch’s t-test. Viral titer data was log transformed prior to analysis. Visual inspections of the transformed data as well as the model’s residuals were examined to ensure the assumptions of normality were met. Corrections for multiple comparisons were conducted using the Holm method when appropriate.

## RESULTS

### Generation of recombinant BCG

SARS-CoV-2 USA-WA1/2020 (passage 4 stock) was obtained from BEI resources (Catalog number NR-52281) and passaged in Vero E6 cells to generate passage 6 virus stock as described (39). Viral RNA was generated from the passage 6 virus stock. We generated cDNA using both oligo(DT) and random hexamer primers. SARS-CoV-2 Nucleocapsid (N) and the RBD was amplified by PCR and cloned into pMV306hsp vector, which has the attachment sequence to integrate itself into the chromosomal *attB* site. This vector does not have the phage derived *xis* gene, so is stable even without antibiotic selection (40). This vector also has a strong mycobacterial heat shock protein 60 (hsp60) promoter and the transgene is cloned in-frame with promoter for constitutive expression (41). We amplified SARS-CoV-2 RBD and N genes using gene specific primers that also contained optimized mycobacterium Shine-Dalgarno sequence (41, 42) and sequences overlapping with pMV306hsp vector. RBD and N gene PCR products were then cloned into pMV306hsp vector using NEB HiFi DNA assembly approach. rBCG carrying transgene was generated by electroporation of competent BCG Danish 1331 strain cells and selected on Middlebrook 7H11 agar with kanamycin. Colony PCR using the same primer set used for amplification of gene-specific product was used for identification of rBCG carrying transgene. rBCG was grown in Middlebrook 7H9 media containing kanamycin (20 μg/ml) and expression of transgene in rBCG lysate was confirmed by western blot (Fig 1). We generated two versions of rBCG carrying gene for RBD; one expressing amino acids 319 to 541 (rBCG-RBD-1) and the other expressing amino acids 319 to 591 (rBCG-RBD-2). Importantly, both rBCG-RBD-1 and rBCG-RBD-2 expressed the protein of interest (Fig 1 A). We also confirmed the expression of N protein from rBCG-N clones (Fig 1B).

**Fig 1:**
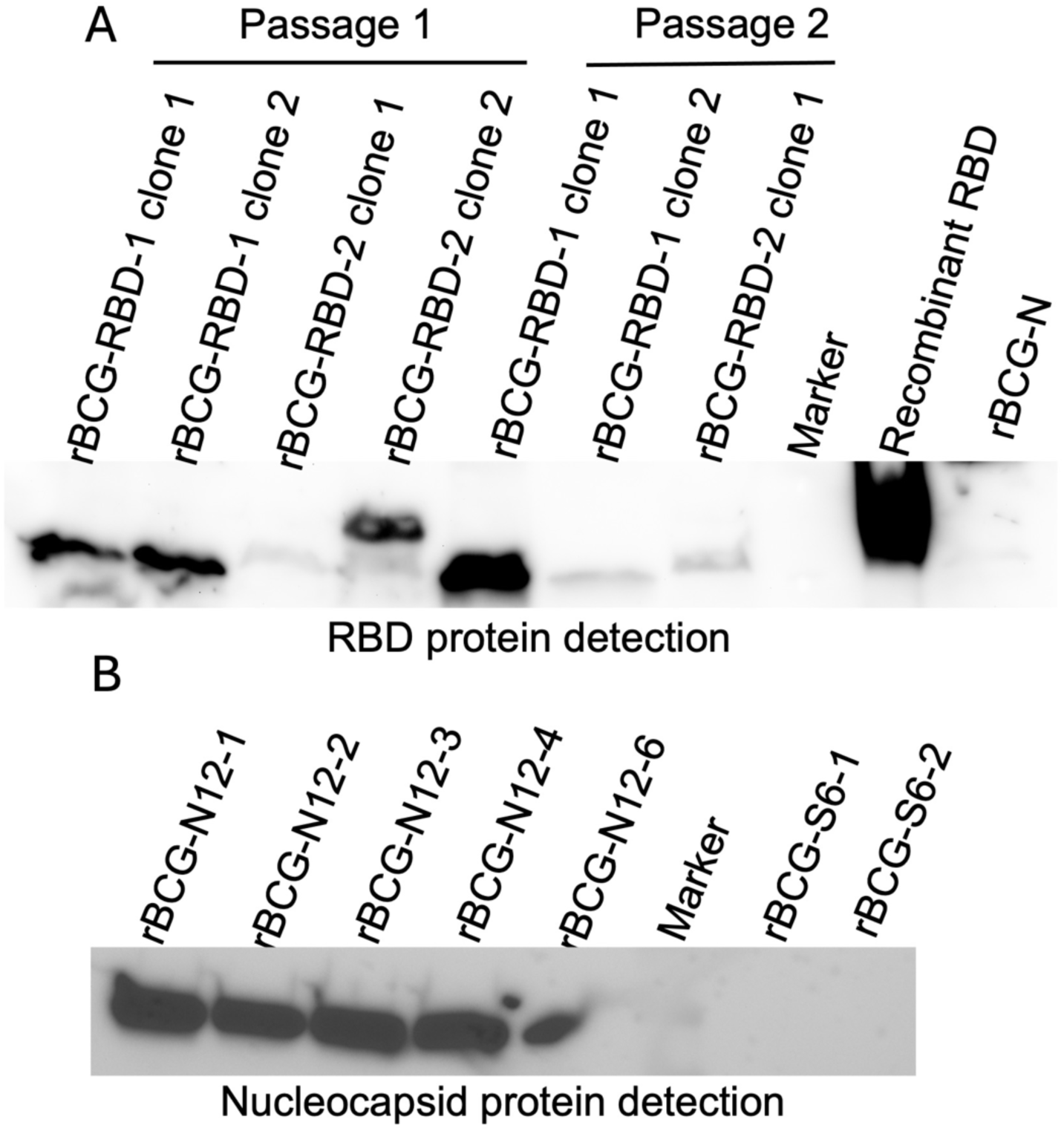
rBCG carrying gene for RBD and N protein of SARS-CoV-2 express protein of interest. A) rBCG-RBD-1 and rBCG-RBD-2 clones 1 and 2 were grown in 7H9 media in the presence of Kanamycin. Bacterial pellet was washed twice with PBS, resuspended in 0.5 to 1 ml of PBS containing protease inhibitor, and transferred to lysing matrix blue tubes (MP). Lysate was prepared using mini bead beater and analyzed by western blot using antibody to N and RBD protein. rBCG-RBD-1 clone 1 and 2 and rBCG-RBD-2 clone 2 show expression of protein of interest in the lysate (A). rBCG-N was used as a negative control. rBCG-RBD-1 clone 1 and rBCG-RBD-2 clone 2 were considered for vaccine preparation. B) rBCG-N clones 1, 2, 3, 4, and 6 were grown in 7H9 media in the presence of Kanamycin. All clones show expression N protein. rBCG clones carrying spike protein gene was used as a negative control.

We further determined the in vitro stability of integrated transgene by subculturing rBCG to 7 to 10 times in 7H9 media with or without Kanamycin (K). After each passage (P), crude lysate was prepared from the bacterial pellet using mini BeadBeater. Fig 2 shows N and RBD protein expression from bacterial lysates prepared from different passages, suggesting in vitro stability of the transgene. These results support published results showing that integration of transgenes into the chromosomal *attB* site are stable (40).

**Fig 2:**
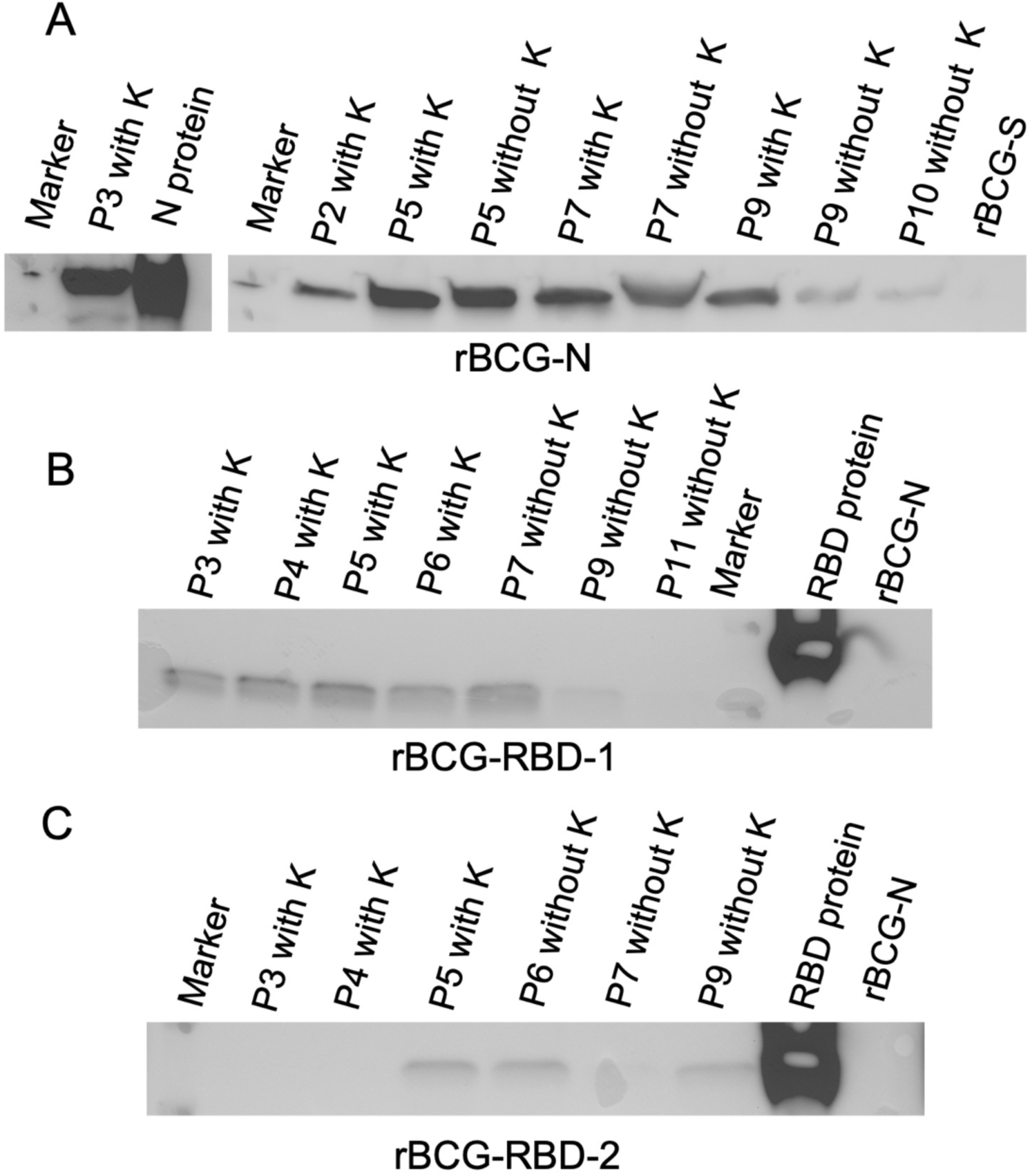
In vitro stability of rBCG carrying transgene of interest. rBCG-N, rBCG-RBD-1, and rBCG-RBD-2 were subcultured in 7H9 media with or without Kanamycin (K) up to 7 to 10 passages. Protein lysate collected at different passage (P) was analyzed by western blot. rBCG-S was used as negative control in Fig 3A, and rBCG-N was used as negative control in 3B and 3C. Recombinant N and RBD proteins were used as positive control. Expression of protein was detected in different passages showing in vitro stability of the transgene in rBCG.

### Immunogenicity and protective efficacy of rBCG prime and protein boost approach

Our initial goal was to develop “prime-boost” vaccination approach, using recombinant BCG expressing SARS-CoV-2 structural proteins for priming and adjuvanted recombinant SARS-CoV-2 structural proteins for boosting. We determined the immunogenicity and protective efficacy of this approach in K18 hACE2 transgenic mouse model, which have been shown to be highly susceptible to SARS-CoV-2 Isolate 2019-nCoV**/**USA-WA1/2020 infection (36). As outlined in Fig 3A, Groups (G) of mice (Table 1) were primed on Day 0 with 1×10^8^ colony forming units of rBCG-RBD-1(G1), rBCG-RBD2 (G2), rBCG-N (G7) or wildtype BCG (G3&G8) in 100 µl of PBS or PBS alone (G4&G9) by subcutaneous injection. At day 42, groups of mice (G1, G2, G3, G4) were boosted with 5 µg of recombinant RBD protein from SARS-CoV-2 Wuhan-Hu-1 strain (BEI resources, NR-52946) formulated in alhydrogel adjuvant 2% (referred to as alum), by subcutaneous injection. G7, G8, and G9 mice were boosted with 5 µg of recombinant N protein formulated in alum. G5 (unvaccinated) mice were injected with alum adjuvant alone. G6 mice surved as naïve controls and were not challenged. We did not observe RBD-specific or N-protein specific antibody response in the serum collected on days 14 and 28, suggesting that rBCG-RBD or rBCG-N priming did not induce the antibody response. We wondered if priming by rBCG-RBD and rBCG-N was below the detection limit and if immunization with alum adjuvanted RBD or N protein would boost the primed antibody response. We measured end-point IgG antibody titer in the serum at Day 56 (14 days post-boosting). Fig 3B shows that RBD protein boost induces antibody response. However, there was no significant difference in the antibody titer between mice primed with rBCG-RBD, wildtype BCG, or PBS alone. Similaly, recombinant N protein boosting induced antibody response. However, there was no significant difference in the antibody titer between rBCG-N, WT-BCG, or PBS primed group. This suggested that rBCG-RBD and rBCG-N did not prime the immune system. Thus, the antibody response observed in the vaccinated mice is mainly due to immunization with recombinant proteins.

**Fig 3.**
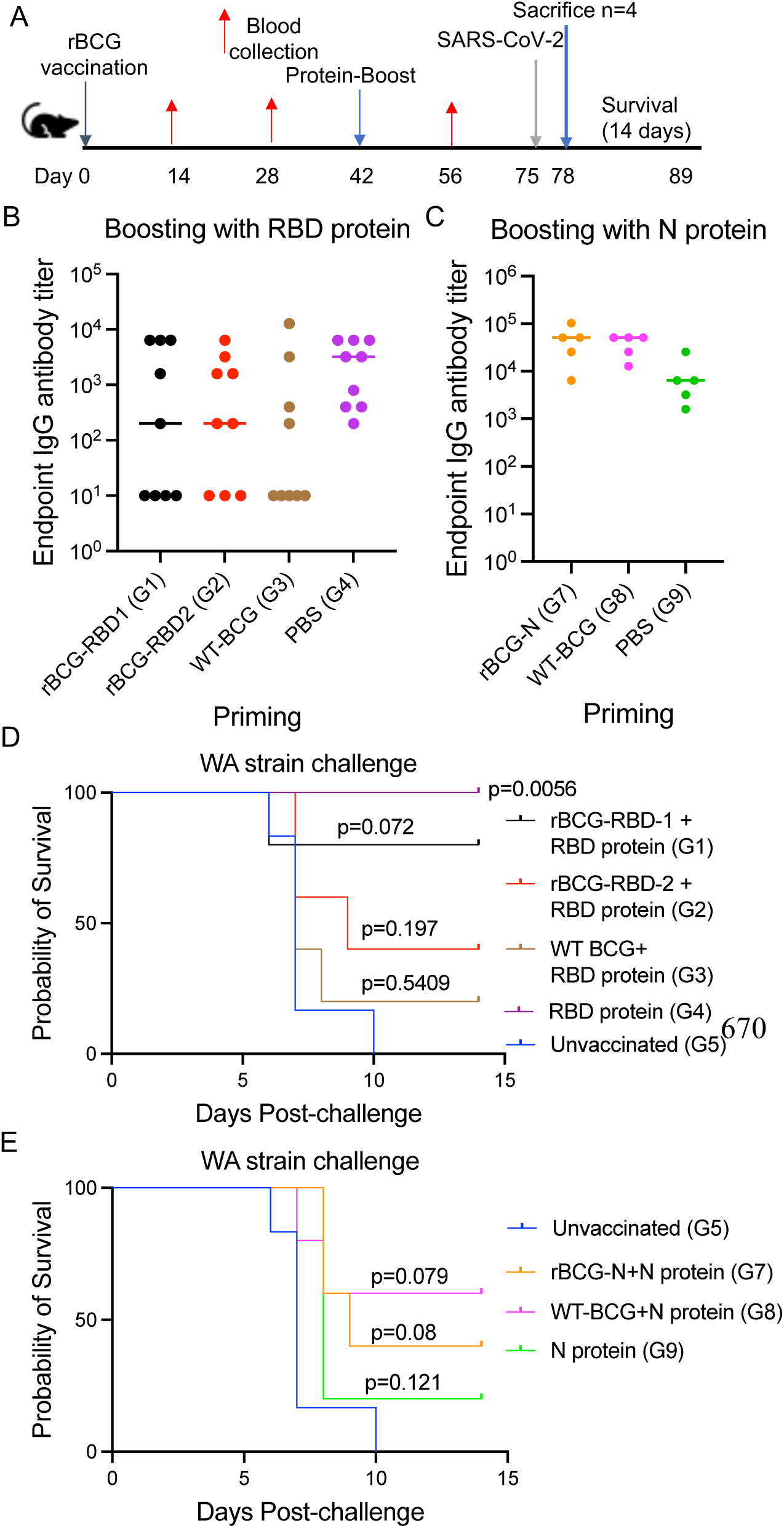
To determine the protective efficacy of rBGG prime and protein boost approach. A) Experimental outline of the study. Mice were primed with rBCG on day 0 and boosted with alum adjuvanted recombinant protein on day 42 by subcutaneous route. Mice were challenged intranasally on day 75 with SARS-CoV-2 WA strain. B&C) RBD and N protein specific binding antibody titer in serum samples collected at day 14 post-boosting, D&E) Kaplan-Meier survival curve analysis between different groups of mice.

**Table 1.** Description of groups of mice used in the study.

| Table 1. Description of Groups of mice used in the study |  |  |  |
| --- | --- | --- | --- |
| Groups | Priming | Boost | Challenge virus |
| 1 (n=9) | rBCG-RBD-1 | Alum<br>adjuvanted<br>RBD protein | SARS-CoV-2<br>WA |
| 2 (n=9) | rBCG-RBD-2 |  |  |
| 3 (n=9) | WT-BCG |  |  |
| 4 (n=10) | PBS |  |  |
| 5 (n=9) | PBS | adjuvant |  |
| 6 (n=9) | PBS | PBS | PBS |
| 7 (n=5) | rBCG-N | Alum<br>adjuvanted<br>N protein | SARS-CoV-2<br>WA |
| 8 (n=5) | WT-BCG |  |  |
| 9 (n=5) | PBS |  |  |

We determined if the antibody response to RBD and N protein provide protection against homologous challenge with WA strain. At day 75 (33 days post-boosting), mice were challenged intranasally with 1×10^3^ pfu of WA strain and followed up for 14 days for survival analysis. Interestingly, mice immunized with RBD protein alone (G4) showed protection against the WA strain. All unvaccinated mice (G5) succumbed to infection by 10 DPC, whereas all RBD protein-vaccinated mice survived to 14 DPC (Fig 1C and supplementary figure 1). Kaplan-Meier analysis showed a significant difference between RBD vaccinated (G4) and unvaccinated (G5) survival curves (p=0.0056, Log-Rank (Mantel-Cox) test). The other group that showed best protection was G1 which received rBCG-RBD-1 prime and RBD protein boost. 4 out of 5 mice (80 % survival) survived the challenge in this group. In rBCG-RBD2 (G2) and WT-BCG (G3) primed group, which received RBD protein boosts, it was not clear why protection against WA strain was not observed. None of the groups immunized with N protein showed protection against SARS-CoV-2 strain. Overall, data suggest that adjuvanted RBD protein from Wuhan strain provides protection against homologous challenge with SARS-CoV-2 WA strain.

### Immunization with recombinant Wuhan RBD protein does not provide protection against heterologous challenge with Delta variant

Intrigued by the above experiment results, we decided to confirm if a single dose alum adjuvanted RBD protein would provide protection against homologous challenge with WA strain and determine if it would provide protection against heterologous challenge with the SARS-CoV-2 B.1.617.2 variant (referred to as Delta variant). As outlined in Fig 4A and Table 2, mice from G1, G3, and G5 were immunized with alum adjuvanted RBD protein by the subcutaneous route. G2, G4, and G6, which received only adjuvant were considered as unvaccinated controls for virus challenge. G7 mice served as naïve controls. Fig 4B shows end-point serum IgG antibody titers in different groups at 14 days post-vaccination (DPV). G1, G3, and G5 mice, which were immunized with alum adjuvanted RBD protein, show comparable anti-RBD specific IgG antibody titer. At 31 DPV, mice were intranasally challenged with 1×10^3^ pfu of WA strain (G1, G2, and G5) or Delta variant (G3 and G4). G6 and G7 were mock-challenged with PBS. Four mice from Groups 1 to 5 were sacrificed at 3 DPC to measure viral loads in lung tissues. 6 mice from Groups 1 to 4 and 11 mice from G5 were followed up to 14 DPC to perform survival analysis. Interestingly, 4 out of 6 mice from G1 and all 11 mice from G5, which were vaccinated with RBD protein, survived the WA strain challenge. Expectedly, all 6 unvaccinated mice (G2) succumbed to WA strain infection (Figure 4C and supplementary figure 2A). Figure 4C showed significant difference in survival curves between G1 v/s G2 (p=0.0048) and G5 v/s G2 (p=<0.0001) survival curves. Furthermore, we also observed significant difference in infectious viral loads (p=0.003, Welch’s t-test) and viral RNA copies (p=0.0011, Welch’s t-test) between vaccinated (4 mice each from G1 and G5) and unvaccinated controls (G2) (Fig 4E&F). Overall, these data confirm that immunization with single-dose alum adjuvanted RBD protein provides protection against homologous challenge with the WA strain.

**Fig 4.**
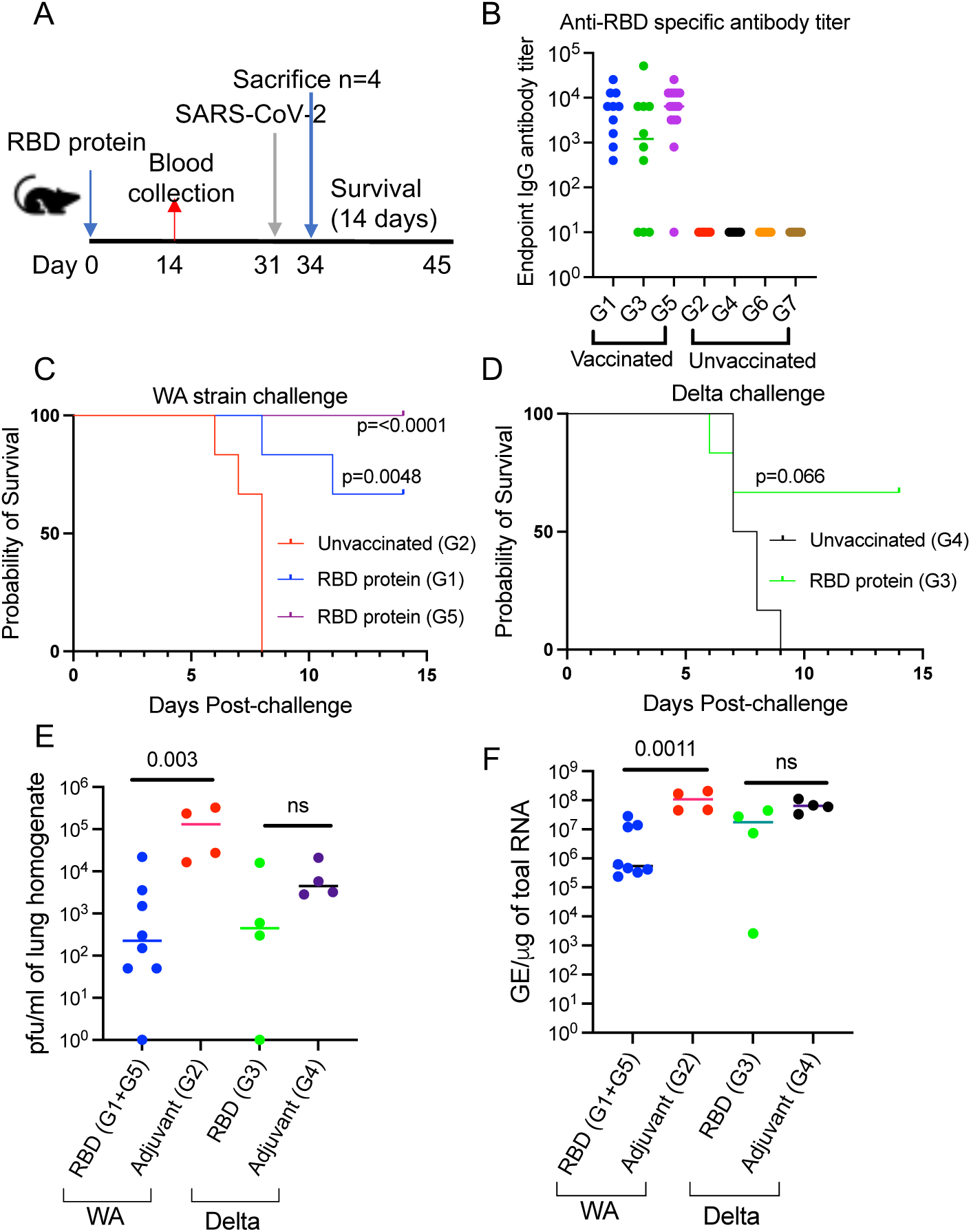
To determine the protective efficacy of single-dose alum adjuvanted RBD vaccine against WA strain and Delta variant challenge. A) Experimental outline. B) RBD-specific binding antibody titer. C) Kaplan-Meier survival curve analysis between different groups of mice following the WA strain challenge. C) Kaplan-Meier survival curve analysis between G3 and G4 mice following Delta variant challenge, E) Infectious viral loads in lung tissues at day 3 measured by plaque assay, F) viral RNA copies in lung tissues at day 3 measured by RT-PCR assay

**Table 2.**
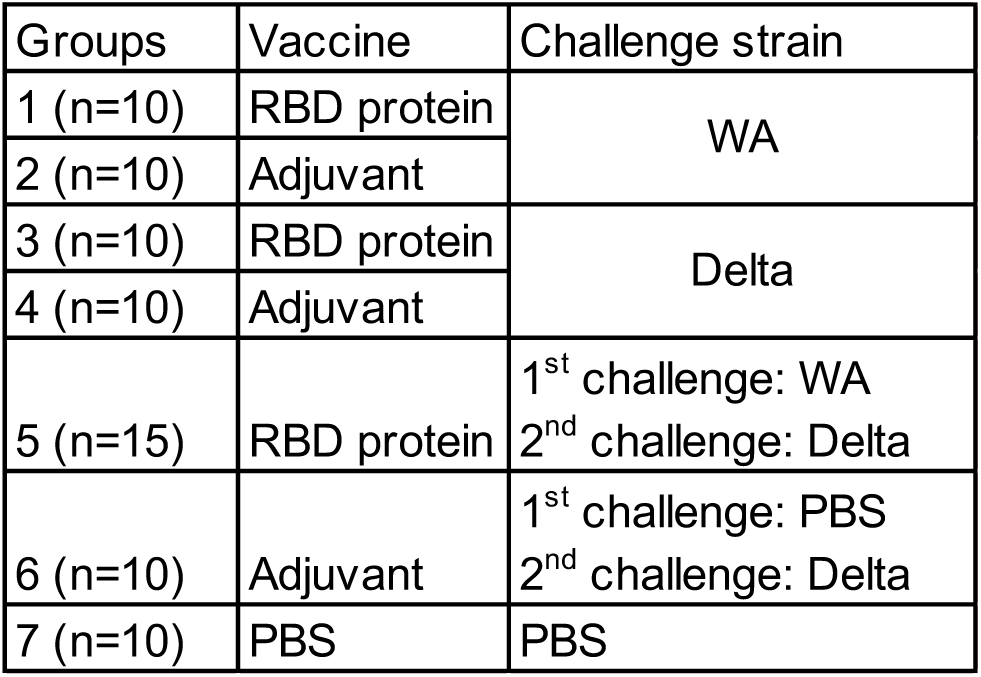
Description of groups of mice used in the study.

| Table 2. Description of groups of mice used in the study |  |  |
| --- | --- | --- |
| Groups | Vaccine | Challenge strain |
| 1 (n=10) | RBD protein | WA |
| 2 (n=10) | Adjuvant |  |
| 3 (n=10) | RBD protein | Delta |
| 4 (n=10) | Adjuvant |  |
| 5 (n=15) | RBD protein | 1<sup>st</sup> challenge: WA<br>2<sup>nd</sup> challenge: Delta |
| 6 (n=10) | Adjuvant | 1<sup>st</sup> challenge: PBS<br>2<sup>nd</sup> challenge: Delta |
| 7 (n=10) | PBS | PBS |

Next, we determined if immunization with single dose RBD protein would provide protection against heterologous challenge with the Delta variant. Interestingly, 4 out of 6 RBD vaccinated mice (G3) survived Delta variant challenge, whereas all 6 unvaccinated mice (G4) succumbed to infection, suggesting approximately 66% of protection against the heterologous challenge (Supplementary figure 2B). However, there was no significant difference in the survival curves between G3 and G4 mice (Fig 4D). Although viral loads in G3 vaccinated mice were low compared to G4 unvaccinated mice, it was not statistically significant (Fig 4E). Similar trend was observed when we measured viral RNA copies of Delta variant by qRT-PCR (Fig 4F). Data suggests that single dose RBD vaccine does not provide protection against heterologous Delta variant challenge.

### Immunization with recombinant RBD protein provides protection against heterologous rechallenge with Delta variant

Many of the current vaccine platforms require at least two doses of vaccines plus the updated booster vaccine. However, this is probably going to be a challenge in resource-limited settings. Along with vaccine hesitancy, there is also going to be the challenge of booster hesitancy. Many individuals vaccinated with either ancestral or recent COVID-19 vaccines are not going to take updated boosters and they have probably been exposed to one or more of SARS-CoV-2 variants. Therefore, it is important to determine if ancestral or recent COVID-19 vaccines combined with virus exposure induces protective immune responses against future variants of concern (VoCs). We hypothesised if RBD-vaccinated mice that survived homologous WA challenge would survive heterologous rechallenge with the Delta variant (Fig 4A). Thus, we rechallenged all G5 vaccinated mice (n=11) with the Delta variant at 14 DPC. G6 mice (n=10) served as unvaccinated controls for Delta variant challenge. At 3 days post-rechallenge (DPrC), 4 mice from G5 and G6 were sacrificed to measure viral loads in lung tissues. Remaining animals were followed for survival. All G5 mice (n=7) survived up to 14 DPrC, whereas the majority of G6 mice (5 of 6) succumbed to Delta rechallenge (Supplementary figure 2C). Fig 5B shows significant difference in survival curves between G5 and G6 mice following Delta rechallenge (p=0.0024). Surprisingly, none of the G5 mice showed detectable infectious virus at 3 DPrC, whereas G6 mice had significantly higher viral loads (p=0.0033, Welch’s t-test) (Fig 5C). Vaccinated mice (G5) also showed significantly lower Viral RNA copy loads compared to unvaccinated controls (G6) Delta rechallenge (p=0.0083, Welch’s t-test) (Fig 5D). Data suggest that single dose RBD vaccine and survival from homologous challenge provides protection against heterologous Delta variant rechallenge.

**Fig 5.**
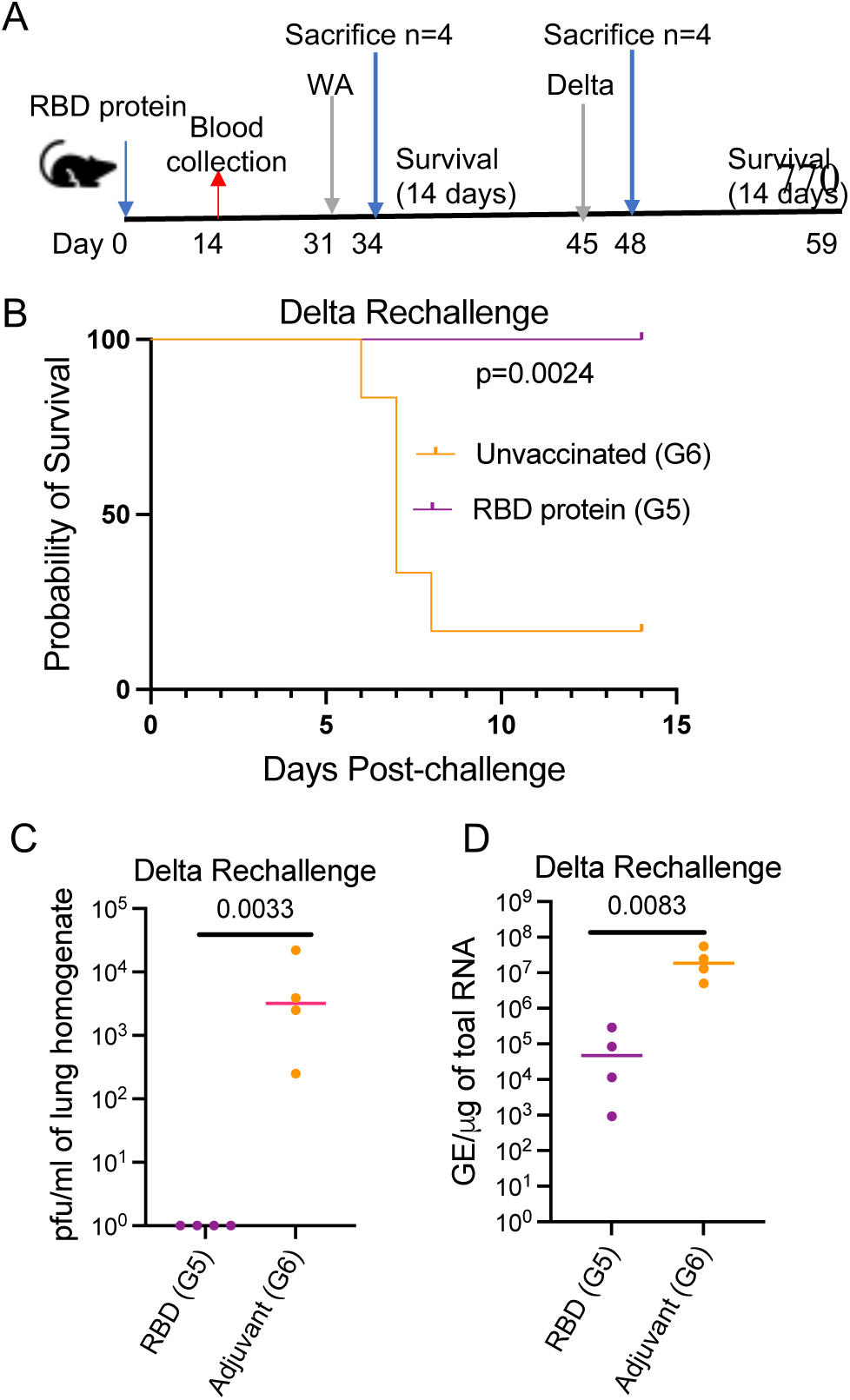
To determine the protective efficacy of single-dose alum adjuvanted RBD vaccine against SARS-CoV-2 rechallenge. A) Experimental outline. B) Kaplan-Meier survival curve analysis between different groups of mice following delta variant rechallenge. C) infectious viral loads of delta variant in lung tissues at 3 day post-rechallenge, D) viral RNA copies in lung tissues at day 3 post-rechallenge measured by RT-PCR assay

### Immunization with recombinant RBD protein provides protection against heterologous rechallenge with Omicron BA.5 variant

Next, we determined if immunization with single dose RBD protein from the ancestral strain provides protection against heterologus challenge and rechallenge with omicron BA.5 variant with an experimental plan very similar to the one performed for Delta variant (Fig 6A&B). Fig 6C shows end-point serum IgG antibody titers in different groups at 14 days post-vaccination (DPV). The omicron BA.5 variant (1×10^5^ pfu/mice) did not cause lethality in K18 hACE2 transgenic mice. Therefore, there was no significant difference in the survival curves between vaccinated and unvaccinated group following BA.5 challenge. We also did not observe significant difference in lung viral loads at 3 DPC (Fig 6D&E), suggesting that immunization with ancestral RBD protein did not protect against BA.5 variant. This also suggests that immunization with Omicron variant specific RBD may be needed for protection against Omicron variant challenge. Interestingly, vaccinated mice that survived WA challenge (n=5) showed undetectable infectious omicron BA.5 viral loads (p=<0.0001) and viral RNA copies (p=<0.0001) in their lungs *vs.* unvaccinated controls (n=10) at 3 DPrC (Fig 6F&G). This again suggests that single dose RBD protein vaccine and survival from WA challenge provides protection against heterologous omicron BA.5 rechallenge.

**Fig 6.**
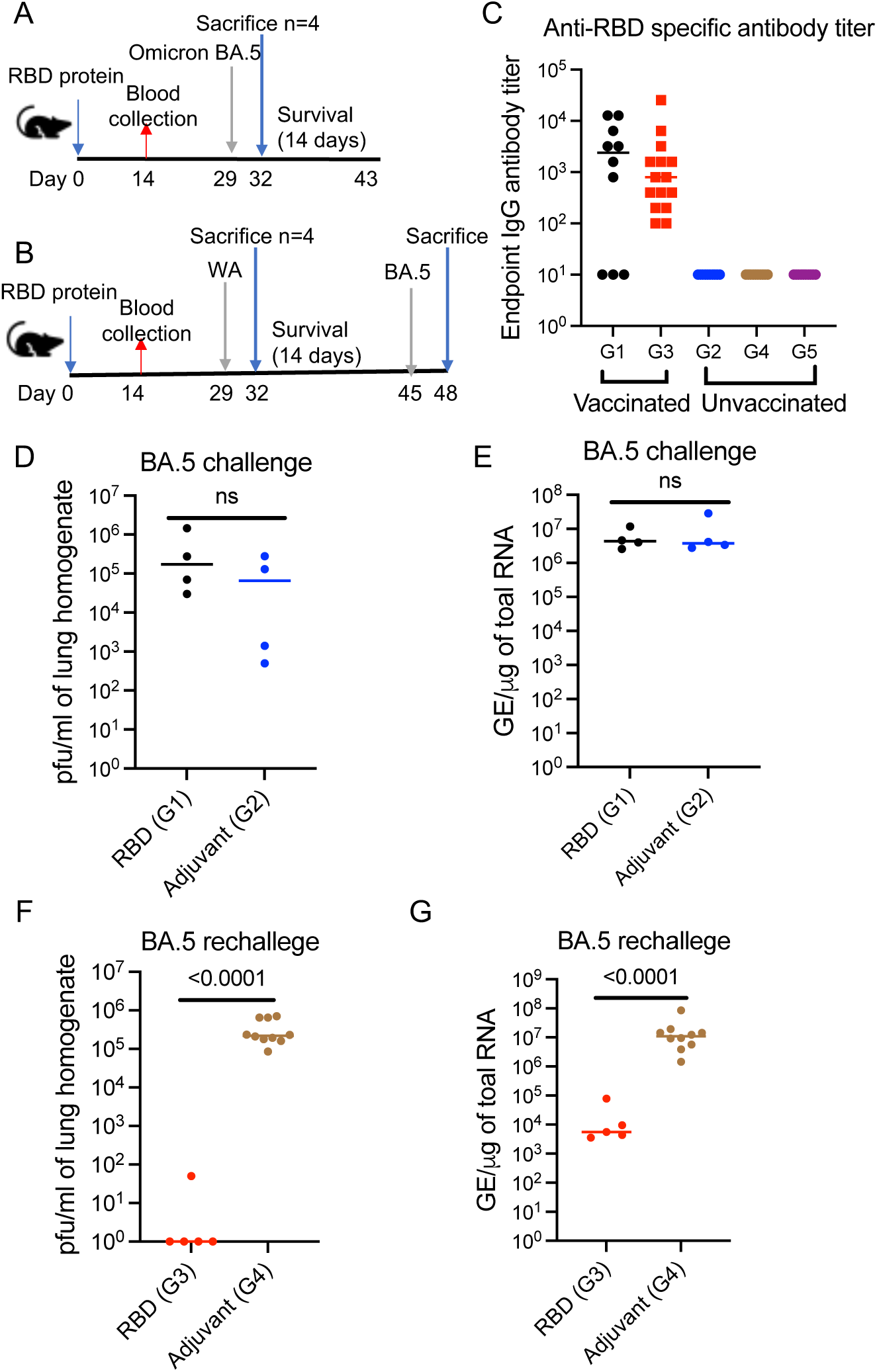
To determine the protective efficacy of single-dose alum adjuvanted RBD vaccine against SARS-CoV-2 challenge and rechallenge. A&B) Experimental outline to determine vaccine efficacy against Omicron challenge (A) and rechallenge (B). C&D) infectious viral loads (C) and viral RNA copies (D) of BA.5 in lung tissues following challenge, E&F) Infectious viral loads (E) and viral RNA copies (F) of BA.5 in lung tissues following rechallenge

### Neutralizing antibody responses in RBD protein vaccinated mice following virus challenge

Neutralizing antibody (NAb) titer in serum samples collected from RBD protein vaccinated mice following virus challenge was determined using a plaque reduction microneutralization assay (37). Neutralization activity is expressed as the 50% inhibitory concentration (IC_50_), which is defined as the reciprocal serum dilution that results in a 50% reduction in pfu compared to virus only control wells. We tested the serum samples collected from groups of RBD vaccinated mice in Table 2 for their ability to neutralize WA strain. Table 4 shows the NAb titers in serum samples collected at different time points post WA and Delta variant challenge. Interestingly, NAb titer was low or below the detection limit at 3 DPC (31 DPV) following WA and Delta virus challenge. However, serum samples collected at 14 DPC showed increase in NAb titer. NAb titer was also observed in serum samples collected after rechallenge with Delta variant. We also tested the serum samples collected from groups of vaccinated mice used in Table 3 for their ability to neutralize the Omicron BA.5 variant. Interestingly neutralizing antibody response against BA.5 variant was observed in serum samples collected at 14 DPC from RBD vaccinated and unvaccinated mice following omicron BA.5 challenge (Table 5). Surprisingly, we could not detect neutralizing antibody response to omicron variant in serum samples collected at Day 3 post rechallenge with omicron variant in RBD vaccinated mice that survived WA challenge (Table 5). Overall, data suggests that RBD protein vaccinated, and virus exposed mice develop neutralizing antibody responses.

**Table 3.** Description of groups of mice used in the study.

| Table 3. Description of groups of mice used in the study |  |  |
| --- | --- | --- |
| 1 (n=10) | RBD protein | Omicron BA.5 |
| 2 (n=10) | Adjuvant |  |
| 3 (n=15) | RBD protein | 1<sup>st</sup> challenge: WA<br>2<sup>nd</sup> challenge: BA.5 |
| 4 (n=10) | Adjuvant | 1<sup>st</sup> challenge: PBS<br>2<sup>nd</sup> challenge: BA.5 |
| 5 (n=10) | PBS | PBS |

**Table 4.** Neutralizing antibody titer against WA strain.

| Table 4. Neutralizing antibody titer against WA strain. |  |  |  |  |
| --- | --- | --- | --- | --- |
| Group description | Mouse ID | 3 dpc | Mouse ID | 14 dpc |
| Group 1: RDB vaccinated and WA challenged | G1-1 | 74 | G1-5 | 954 |
|  | G1-2 | <50 | G1-6 | 1226 |
|  | G1-3 | <50 | G1-7 | 2559 |
|  | G1-4 | <50 | G1-10 | 576 |
| Group 3: RDB vaccinated and Delta challenged | G3-1 | 73 | G3-5 | 2190 |
|  | G3-2 | <50 | G3-7 | 2010 |
|  | G3-3 | 51 | G3-8 | 8303 |
|  | G3-4 | <50 | G3-10 | 2000 |
| Group 5: RDB vaccinated and WA challenged | G5-1 | <50 |  |  |
|  | G5-2 | <50 |  |  |
|  | G5-3 | 109 |  |  |
|  | G5-4 | <50 |  |  |
|  |  | 17 dpc (WA)/<br>3DPrC (Delta) |  | 28 dpc (WA)/<br>14DPrC (Delta) |
| Group 5: RDB vaccinated, WA challenged and Delta rechallenged | G5-5 | 6213 | G5-9 | 533 |
|  | G5-6 | 571 | G5-10 | 1134 |
|  | G5-7 | 93 | G5-11 | 5114 |
|  | G5-8 | 418 | G5-12 | 1983 |
|  |  |  | G5-13 | <50 |
|  |  |  | G5-14 | <50 |
|  |  |  | G5-15 | <50 |

\*Serum samples from RBD vaccinated mice following virus challenge (see Table 2) were tested.

**Table 5.** Neutralizing antibody titer against BA.5 variant.

| Table 5. Neutralizing antibody titer against BA.5 |  |  |
| --- | --- | --- |
| Group description | Mouse ID | 14 dpc |
| Group 1: RDB vaccinated and BA.5 challenged | G1-5 | 3238 |
|  | G1-6 | 8794 |
|  | G1-7 | <100 |
|  | G1-8 | 3698 |
|  | G1-9 | <100 |
|  | G1-10 | 3608 |
| Group 2: unvaccinated and BA.5 challenged | G2-5 | 2708 |
|  | G2-6 | 4435 |
|  | G2-7 | 4286 |
|  | G2-8 | 2820 |
|  | G2-9 | 3807 |
|  | G2-10 | 4494 |
|  | Mouse ID | 17 dpc (WA)/<br>3DPrC (BA.5) |
| Group 3: RDB vaccinated, WA challenged and BA.5 rechallenged | G3-5 | 141.9 |
|  | G3-7 | 66.96 |
|  | G3-9 | <50 |
|  | G3-13 | <50 |
|  | G3-15 | <50 |

\*Serum samples from RBD vaccinated and unvaccinated mice following virus challenge (see Table 3) were tested.

## DISCUSSION

Currently, there are 12 WHO-approved COVID-19 vaccines granted for emergency use or full approval. These include mRNA-based vaccines (Pfizer and Moderna), nonreplicating adenovirus-based vaccines (Jansen and Oxford/AstraZeneca), protein subunit vaccines based on full-length spike protein (Novavax), and inactivated SARS-CoV-2 vaccines (Bharat Biotech, and Sinopharm). These vaccines have played a significant role in addressing the pandemic. However, the emergence of new SARS-CoV-2 variants of interest and variants of concern (VoCs) with increased transmissibility and antibody resistance (43–47), as well as vaccine breakthrough infections (48–52), have complicated the control of the COVID-19 pandemic. Furthermore, waning immunity and the continued evolution of the virus has also necessitated the development of variant-specific or pan-coronavirus-specific vaccines. Therefore, there is still a great need for developing novel vaccines that can better prevent SARS-CoV-2 infection, transmission, and viral replication.

Our initial goal was to develop rBCG prime and protein boost approach as a vaccine strategy for COVID-19. Interestingly, BCG vaccination has also been shown to have non-specific beneficial effects against unrelated pathogens (31). It has been hypothesized that the non-specific beneficial effects of BCG vaccination are due to long-term activation and reprogramming of innate immune cell memory, a phenomenon termed trained immunity (32–34). When activated by unrelated infection, these epigenetically modified cells respond rapidly resulting in enhanced protective immune response (33, 34). Establishment of trained immunity by BCG vaccination has led to the hypothesis that that it might provide protection against severe form of COVID-19 disease (30, 35, 53, 54). Establishment of trained immunity by BCG vaccination has led to the hypothesis that it might provide protection against severe form of COVID-19 disease (30, 35, 53, 54). Recent epidemiological studies suggested possible association between BCG vaccination and reduced risk of COVID-19 (55–58). Although conflicting reports exist (59, 60), some clinical trials suggest that BCG vaccination may provide some protection against COVID-19 (61–64). In a mouse study, single dose, BCG-adjuvanted stabilized, trimeric form of the spike protein vaccine provided protection against SARS-CoV-2 infection in K18 hACE2 mice (65). In another study, intravenous, but not subcutaneous, administration of BCG completely protected mice against lethal SARS-CoV-2 challenge (66). These advantages of BCG vaccination led us to hypothesize that rBCG expressing SARS-CoV-2 structural genes will induce both specific and non-specific beneficial immune responses, thereby providing better protection against SARS-CoV-2 infection. rBCG expressing antigens from various pathogens to induce both humoral and cell-mediate immunity have been developed (67). In the case of respiratory tract infections, rBCG expressing respiratory syncytial virus (RSV) and human metapneumovirus (hMPV) proteins have been shown to be immunogenic and protective against respective pathogen challenges in mouse models (68, 69). Therefore, we have generated rBCG carrying SARS-CoV-2 structural genes such as nucleocapsid and the RBD of spike protein, which are immunological targets for vaccine development (70–74). However, subcutaneous inoculation of rBCG-RBD and rBCG-N did not induce antibody responses to RBD or N proteins, respectively. It is not clear why these rBCG clones were not immunogenic in mouse models. We speculate that it could be due to poor expression of the transgenes in mice. We also determined the protective efficacy of rBCG prime and protein boost approach in transgenic mouse model against SARS-CoV-2 challenge. However, rBCG priming did not enhance the protection afforded by protein immunization, as the RBD protein only vaccinated group showed complete protection against WA strain compared to rBCG-RBD or WT-BCG primed and RBD protein boosted groups. It is also not clear, why the RBD protein vaccine did not show protection against WA strain in mice that received rBCG or WT BCG prime.

It is interesting that a single dose alum adjuvanted RBD protein vaccine from ancestral Wuhan strain provides protection against homologous challenge. Expectedly, single dose RBD vaccine from ancestral Wuhan strain did not provide protection against heterologous challenge with SARS-CoV-2 Delta and Omicron variants. Perhaps two doses of RBD vaccine could have provided protection. Lack of protection against VoCs could be likely due to the ability of these variants to overcome neutralizing antibody responses generated by the RBD protein of the Wuhan strain. This also suggests that variant specific or pan-beta-CoV vaccines may be needed to see protection against variants of concern. However, it takes time and may not be cost effective. Therefore, we wondered if RBD protein vaccinated mice that survive homologous challenge with WA strain show protection against VoCs. Interestingly, we observed that RBD vaccinated mice that survived the WA strain challenge showed protection against rechallenge with the Delta and BA.5 VoCs. These findings suggest that a single-dose vaccine combined with survival from virus challenge induces multiple protective immune responses against heterologous challenge with the Delta and BA.5 VoCs. We expect that virus challenge to result in an anamnestic response and enhance the immune responses induced by single dose RBD vaccines. Virus exposure will also induce immune responses to other immunological targets of SARS-CoV-2, such as nucleocapsid, membrane, and envelope proteins, which should contribute to protection against rechallenge with SARS-CoV-2 variants.

Live attenuated vaccines are considered the best to induce multiple immune responses. Interestingly, RBD vaccinated mice showed lower viral loads compared to the unvaccinated mice following WA strain challenge. This suggests that RBD vaccine potentially makes the pathogenic virus into an attenuated strain. Therefore, we propose that a vaccine against a homologous virus may be used as a concept to generate live attenuated vaccine. Our data also supports that vaccine plus virus exposure may induce multiple immune responses providing protection against challenge with SARS-CoV-2 variants.

Many of the currently approved vaccine platforms require at least two doses of primary vaccine followed by updated booster vaccine. However, this is probably going to be a challenge in resource-limited settings. Along with vaccine hesitancy, there is also going to be the challenge of booster hesitancy. Thus, it is important to determine if vaccinated individuals exposed to SARS-CoV-2 infection develop multiple immune responses that may protect them against future variants of concern (VoCs). Our mouse models clearly indicate that vaccine and virus expose will provide protection against Delta and BA.5 variants. it would be interesting to determine if this approach would show protection against more recent SARS-CoV-2 variants.

One caveat with our rechallenge study design is that mice were rechallenged 14 days after the first challenge. Therefore, the residual innate immune responses present after the first virus challenge may also play a role in protection against SARS-CoV-2 challenge. Thus, there is a need to study durability of protective immune responses induced in the context of infection in vaccinated mice. Another caveat in the study design is the absence of a group with virus only exposed mice. We could not have this group in the study as majority of the unvaccinated mice exposed to WA strain succumb to infection.

It is not going to be practical to conduct placebo-controlled vaccine efficacy clinical trials in humans. Thus, understanding the vaccine-induced immune responses that provide protection against SARS-CoV-2 infection will be useful in the design and development of future effective vaccines. We have mouse models where RBD vaccinated mice show protection against WA strain but not against Delta and BA.5 variants. We also have mouse models where RBD vaccinated mice that survive homologous challenge show protection against heterologous rechallenge with Delta and Omicron BA.5 VoCs. Our mouse models of vaccine mediated protection and no protection will be very useful in determining immune correlates of protection. Identifying protective immune responses will be critical for the development of new and updated vaccines against SARS-CoV-2 and other coronaviruses.

Overall, our data suggest that single does RBD vaccine combined with virus exposure provides protection against SARS-CoV-2 variants of concern. In a pandemic setting, it is important to vaccinate the population with at least one dose of the vaccine, which should provide some degree of protection against circulating variant. These vaccinated and virus exposed individuals may develop protective immune responses against future VoCs.

## Author contributions

RT: obtained funding, designed studies, performed experiments, analyzed data, wrote manuscript; VK: designed mouse studies, provided resources, and edited manuscript; VD: designed mouse studies and edited manuscript; JM, MA, JC, OR, and BM: performed experiments; MG: standardized qRT-PCR assay and edited manuscript; RM: performed statistical analysis of the data; DK: Mentored RT, provided resources, and edited manuscript. RC: Mentored RT, provided funding support and resources, and edited manuscript.

## Funding support

Funding support to RT: Texas Biomed Forum Grant-2021, Texas Biomed Forum Grant-2022, 1R21AI165289, Texas Developmental Center for AIDS Research pilot grant (P30AI161943-03), and 1R03AI174973

## Acknowledgments

The following reagent was produced under HHSN272201400008C and obtained through BEI Resources, NIAID, NIH: Spike Glycoprotein Receptor Binding Domain (RBD) from SARS-Related Coronavirus 2, Wuhan-Hu-1 with C-Terminal Histidine Tag, Recombinant from HEK293T Cells, NR-52946.

---

**Supplementary Figure 1:**
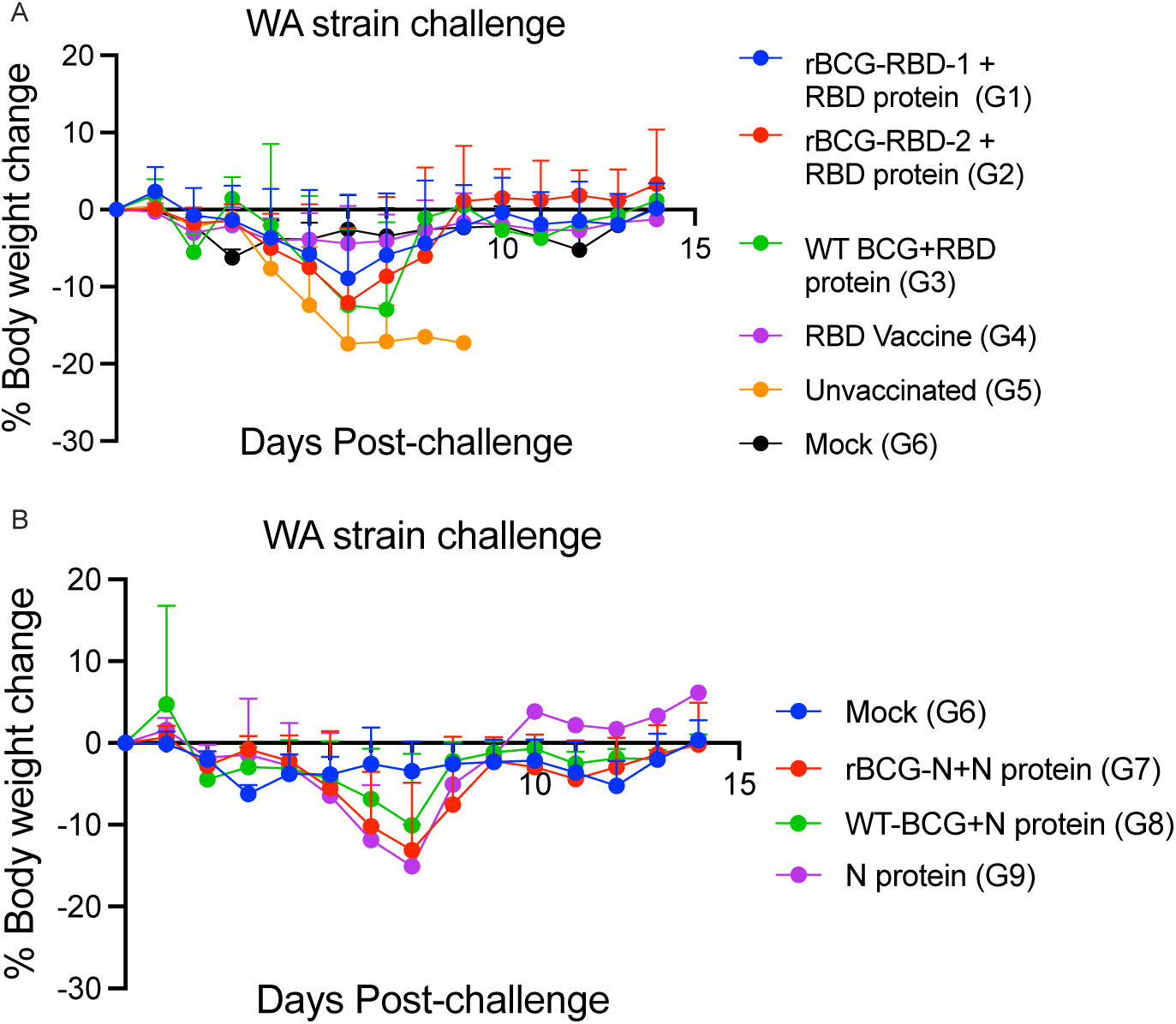
Percentage change in body weight following SARS-CoV-2 WA challenge. Body weights were measured every day post-virus challenge. Baseline body weight is the body weight measured on the day of challenge. Percentage change from baseline for each of the group is depicted in the graph.

**Supplementary Figure 2:**
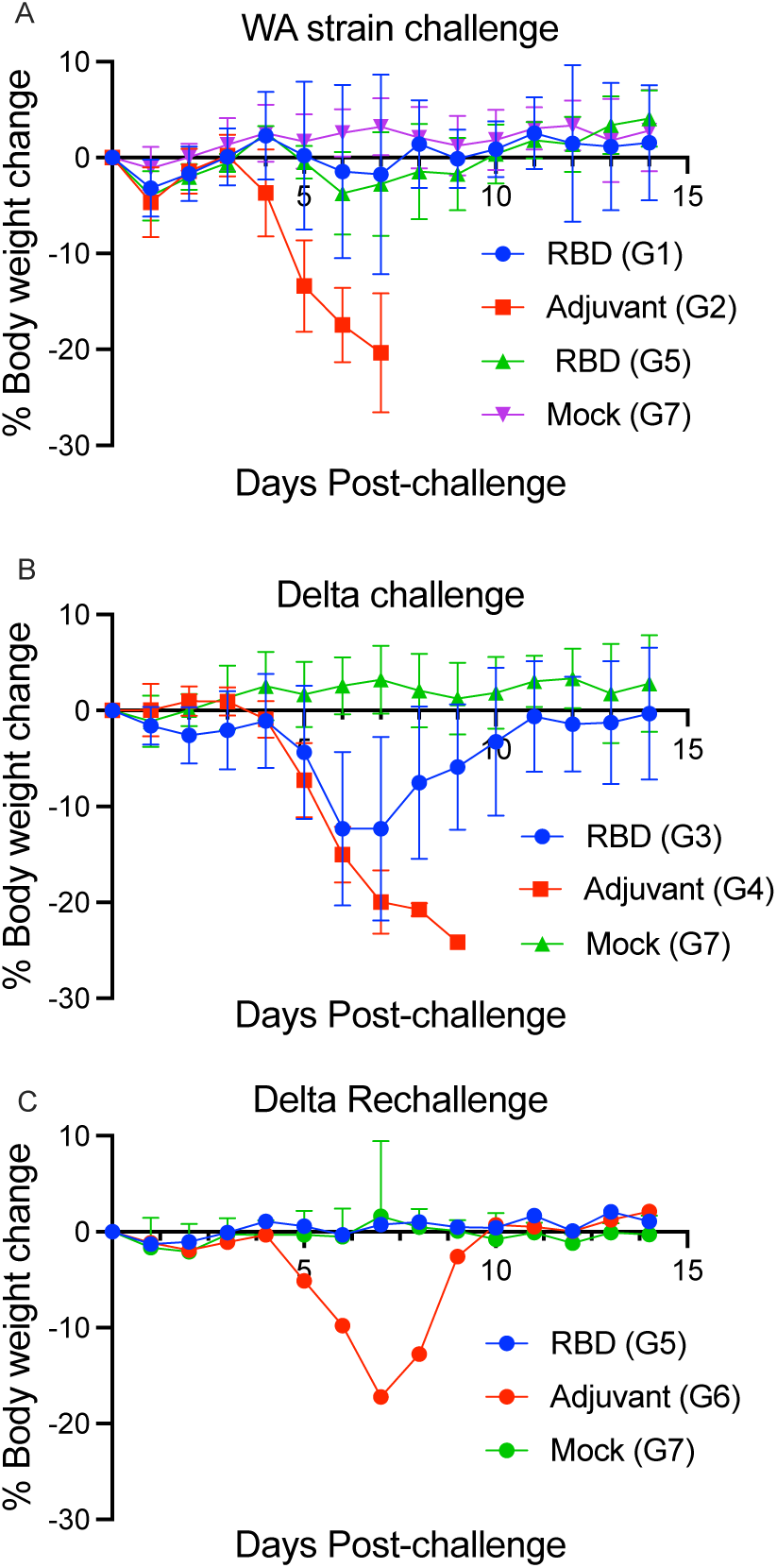
Percentage change in body weight following SARS-CoV-2 challenge. Body weights were measured every day post-virus challenge. Baseline body weight is the body weight measured on the day of challenge. Percentage change from baseline for each of the group is depicted in the graph.

## Notes

### Competing Interest Statement

The authors have declared no competing interest.

